# Genetic Dissection of the role of *Piga* and *Pgap2* in the embryonic mouse brain

**DOI:** 10.64898/2025.12.08.692975

**Authors:** Jennifer L. Watts, Jesus M. Leal, Rolf W. Stottmann

## Abstract

Glycosylphosphatidylinositol (GPI) anchors are a post-translational modification of over 150 proteins. These GPI-anchored proteins are enriched in lipid rafts in the plasma membrane and serve a variety of functions including acting as receptors and cell adhesion molecules. Human pathogenic variants in GPI biosynthesis pathway enzymes are collectively called inherited GPI deficiencies and lead to several brain anomalies such as global developmental delay, hypomyelination, cerebellar hypoplasia, and premature death. PIGA is the first catalytic enzyme in the GPI anchor biosynthesis cascade, which creates the backbone of all GPI anchors, and PGAP2 is one of the last enzymes in the GPI-anchor biosynthesis pathway and modifies the protein-bound anchor for trafficking to the cell membrane. A *Nestin*-*Cre* mediated deletion of *Piga* in the mouse led to early postnatal death and significant structural brain malformations, similar to pathogenic human *PIGA* variants. While this model provides an opportunity to develop treatments for these neurological phenotypes, we still lack a deep understanding of GPI-anchor functions in brain development. Additionally, there are no studies on *Pgap2* in the brain to date. We extended these studies with a series of genetic ablations to precisely determine the role of *Piga* and *Pgap2* in the forebrain, oligodendrocytes, and cerebellum. We find *Piga* expression in the hindbrain is absolutely required for survival while *Pgap2* ablations were much less deleterious. These studies broaden our knowledge on the brain region-specific requirements of GPI-anchor biosynthesis enzymes.

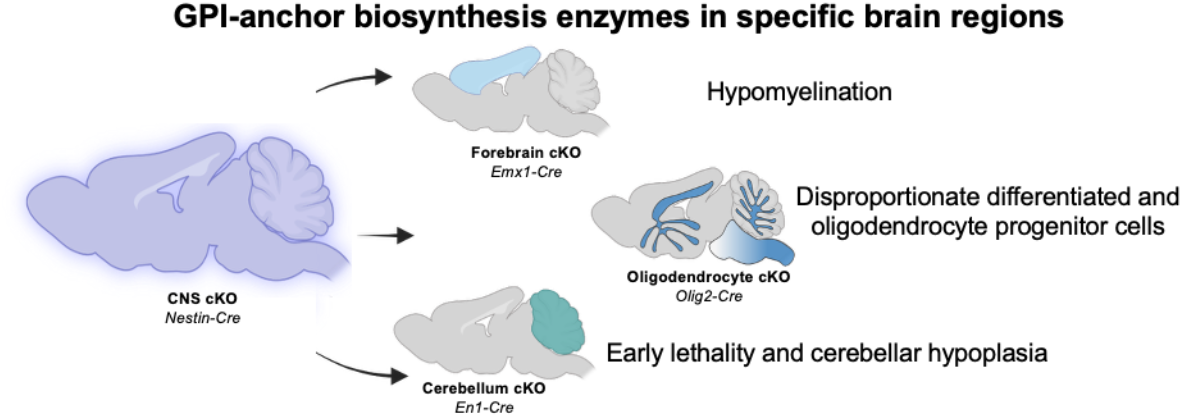

**Significance Statement:** In this study, we examine the roles of the GPI-anchor biosynthesis enzymes *Piga* and *Pgap2* to understand the poorly understood mechanisms underlying brain anomalies seen in the inherited GPI deficiencies. Using Cre/lox conditional mouse genetic models, we deleted *Piga* and *Pgap2* from various brain regions, including the forebrain, oligodendrocytes, and cerebellum, and observed brain abnormalities. We find that *Piga* ablations mimic many brain phenotypes seen in human inherited GPI deficiencies, while *Pgap2* ablations do not grossly disrupt neurodevelopment. Additionally, we demonstrate that *Piga* expression is essential in the cerebellum for survival. These findings reveal mechanisms underlying several inherited GPI deficiency brain phenotypes and will inform targeted therapeutic strategies.

## Introduction

Posttranslational modifications of proteins are often essential for proper function and molecular interactions. Glycosylphosphatidylinositol (GPI) anchors are a class of post-translational modifications that attach to more than 150 proteins localized in the plasma membrane to facilitate function as adhesion molecules, membrane enzymes and receptors (1-3). GPI-anchor biosynthesis utilizes a cascade of twenty-eight enzymes starting with Phosphatidylinositol Glycan Biosynthesis Class (PIG) proteins in the endoplasmic reticulum (2, 4-8). After the GPI-anchor backbone is attached to proteins, the GPI-anchored proteins translocate to the Golgi apparatus. Post-GPI Attachment to Protein (PGAP) enzymes in the Golgi modify fatty acid chains on the GPI-anchor to facilitate the stabilization of the protein bound GPI-anchor at the cell surface (9, 10).

Pathogenic variants in genes encoding these enzymes are linked to numerous congenital disorders, collectively termed Inherited GPI Deficiencies. While there is great variability in disease severity, common neural and craniofacial phenotypes include developmental delay, epilepsy, cleft lip/palate, dysmorphic features, and a significant fraction of these patients die prematurely (11). Some Inherited GPI Deficiency patients show structural brain anomalies including delayed white matter development and/or atrophy, thin corpus callosum, cerebellar hypoplasia and/or atrophy, and microcephaly (12-33). To date, there is no comprehensive understanding of GPI-anchor deficiency disorders and current treatment options are mostly limited to managing symptoms (11, 22, 30, 34). Therefore, studying GPI-anchor biosynthesis throughout development is essential to better treat patients with this growing prevalence of disorders.

Phosphatidylinositol Glycan Biosynthesis Class A (PIGA) is part of the glycosylphosphatidylinositol-N-acetylglucosaminyltransferase complex and initiates GPI-anchor biosynthesis attaching a N-acetylglucosamine to the phosphatidylinositol molecule on the cytoplasmic side of the endoplasmic reticulum. *PIGA* is required for the biosynthesis of virtually all GPI-anchors in cells (35, 36). PIGA is located on the X chromosome and pathogenic variants therefore cause an X-linked recessive disorder. Human female carriers are unaffected due to skewed X-inactivation, consistent with a detrimental effect in cells with inactivation of the wild-type allele (11, 19, 20, 24, 25, 27). Post-GPI Attachment to Proteins 2 (PGAP2) is one of the last GPI-anchor biosynthesis enzymes and modifies the fatty acid chain on the GPI-anchor in the Golgi apparatus (9, 10, 36-38). While PGAP2 is not absolutely required for GPI-anchor development, it does reduce the amount of GPI-anchored protein within the plasma membrane (35, 36). *PGAP2* is an autosomal gene and disease follows a recessive form of inheritance. Even though both enzymes are in the GPI-anchor biosynthesis pathway, *PGAP2* variants often have less severe phenotypes than *PIGA* variants (12, 15, 26, 31, 37, 39). This is likely because previous studies have found the while loss of *PIGA* leads to a complete loss of GPI-anchor proteins, loss of *PGAP2* only has a partial effect (35, 36).

Mouse models have facilitated fundamental discoveries into the pathogenesis of inherited GPI deficiencies. *Piga* and *Pgap2* null mice are both embryonic lethal prior to organogenesis which establishes a need for hypo-morphic or conditional alleles to study GPI-anchor biosynthesis in developing tissues (36, 40). We previously published a mouse model with ablation of *Piga* in the central nervous system (*Nestin-Cre; Piga*) and observed phenotypes including cerebellar hypoplasia with abnormal Purkinje cells, ataxia, hypomyelination, and premature death (41) which are reminiscent of patients with *PIGA* pathogenic variants. We have also previously identified the *Pgap2*^*clpex*^ hypo-morphic allele for *Pgap2* via an ENU mutagenesis screen (36, 42). Though the major phenotypes in these studies involved neural tube and craniofacial defects, we used the conditional gene trap allele *Pgap2*^*tm1a(KOMP)Vlcg*^ (*Pgap2* ^*null*^) to show distinct *Pgap2* forebrain expression in a domain consistent with the location of the intermediate, or basal, progenitors in embryonic day (E)16.5 brains. *PGAP2* patients also have intellectual disability and developmental delay with some cases of microcephaly or macrocephaly.

The *Nestin-Cre* transgene drives Cre recombination activity broadly throughout the brain at neurogenesis stages (E11.25) (43). As this is a fairly broad recombination pattern, we wanted to more specifically understand the consequences for *Piga* deletion from different portions of the brain to potentially identify which region(s) were most affected by deletion of *Piga* and might be causing the lethality seen in the *Nestin-Cre; Piga* model. In this study, we used *Emx1-Cre, Olig2-Cre*, and *En1-Cre* mouse lines to further investigate PIGA function in distinct neuronal tissues and cells such as the forebrain, oligodendrocytes, and the cerebellum (43-46). Additionally, we investigated the requirement for PGAP2 function in the forebrain (*Emx1-Cre*) and the cerebellum (*En1-Cre*) to directly compare two roles for these GPI-anchor biosynthesis enzymes in neurodevelopment.

## Results

### Deletion of Piga in the dorsal forebrain causes mild cortical defects and hypomyelination

One of the structural brain malformations often seen in patients with *PIGA* variants is microcephaly. We previously published a *Nestin*-Cre mouse model, a radial glial-specific conditional knock-out, which did not display a microcephaly phenotype (41). This could possibly be due to the recombination of *Nestin-Cre* in nervous tissue starting at E11.5 which might be too late to cause changes in neuroprogenitors, leading to microcephaly (41, 43, 47). Therefore, we genetically ablated *Piga* in the entire dorsal forebrain using the *Emx1-Cre* (*B6*.*129S2-Emx1*^*tm1(cre)Krj/J*^) with robust recombination evident by E10.25 (44). We mated *Emx1*^*Cre*^ males with *Piga*^*flox/X*^ females to produce both *Emx1*^*Cre*^; *Piga*^*flox/X*^ females and *Emx1*^*Cre*^; *Piga*^*flox/Y*^ males to determine the consequences of ablating *Piga* throughout the dorsal forebrain.

At postnatal day (P) 21, we observed no reduction in the recovery of mutants (n=62, Chi-square at P21 p=0.406, Fig.1A). Interestingly, *Emx1*^*Cre/wt*^; *Piga*^*flox/Y*^ males start to die shortly after weaning (n=8, Mantel-Cox p=0.107) whereas *Emx1*^*Cre/wt*^*;Piga*^*flox/X*^ females can survive up to 45 days post birth (n=6, SFig.1A). *Emx1*^*Cre/wt*^; *Piga*^*flox/Y*^ mutant male body weights (n=5) are significantly lower than wildtype animals (n=25, p<0.001) and *Emx1*^*Cre/wt*^; *Piga*^*flox/X*^ females (n=4, p=0.016) by P22 (SFig.1B). These results reveal that *Piga* expression in the mouse forebrain is required for the animal to thrive after P21.

**Figure 1.**
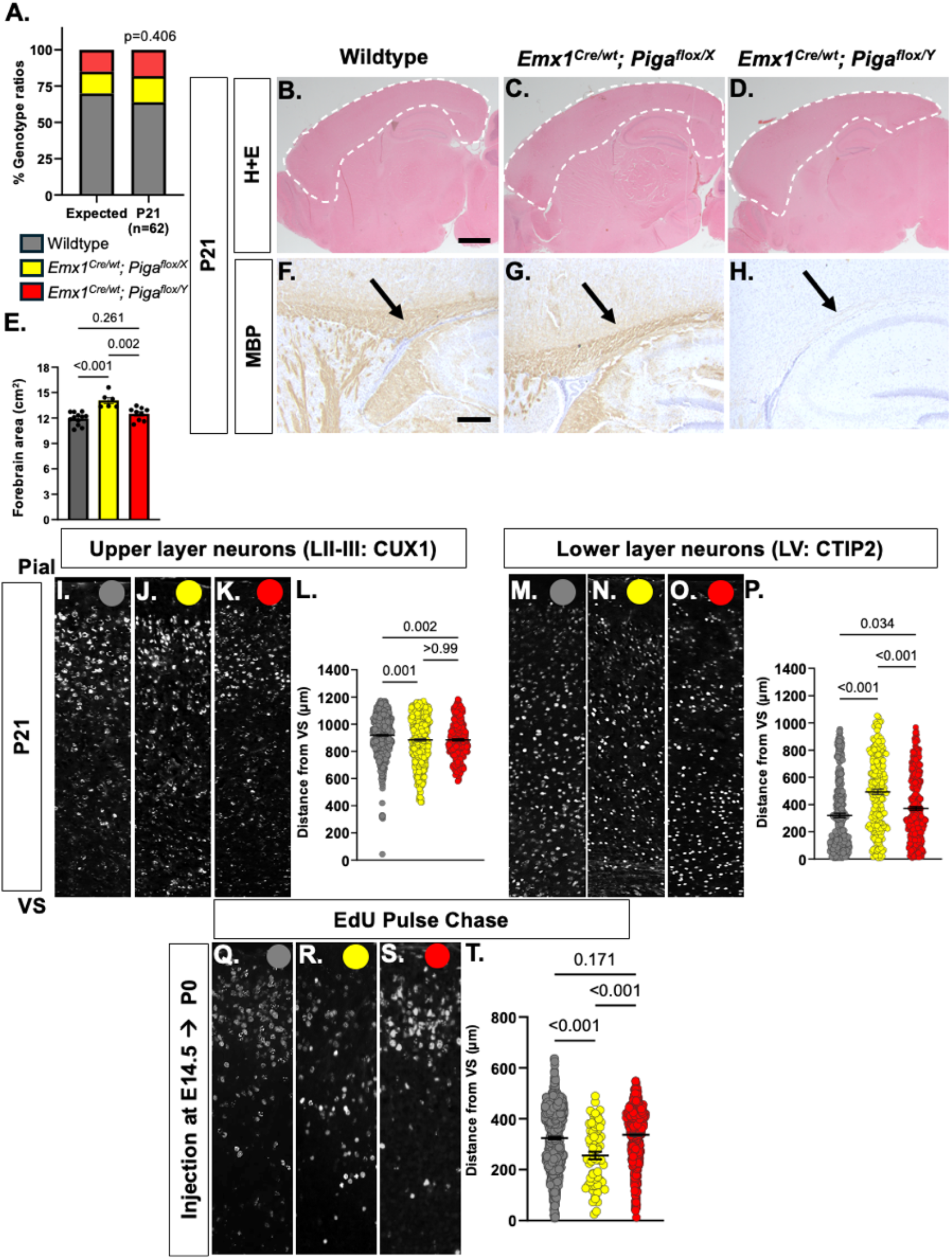
*Piga* depletion from the forebrain causes mild cortical defects and hypomyelination. **A)** Genotype ratios of forebrain ablated *Piga* animals are similar to expected genotype ratios at P21 (n=62, Chi-square p=0.406). Hematoxylin and eosin (H+E) images of wildtype, **C)** *Emx1*^*Cre/wt*^; *Piga*^*flox/X*^ female and **D)** *Emx1*^*Cre/wt*^; *Piga*^*flox/Y*^ male P21 forebrain sections (n=6-12 animals, scale bar= 1 mm). **E)** Quantification of forebrain area in sections highlighted in images B-C (ANOVA: p<0.001). Immunostaining of Myelin Basic Protein (MBP)(**F-H**), CUX1 (**I-K**) and CTIP2 (**M-O)**, and EdU (**Q-S**) in wildtype (**F, I, M, Q)**, *Emx1*^*Cre/wt*^; *Piga*^*flox/X*^ female (**G, J, N, R**), and *Emx1*^*Cre/wt*^; *Piga*^*flox/Y*^ male (**H, K, O, S**) P21 forebrain sections (n=4, scale bar= 250 μm). Quantification of CUX1 (**L**, n=4 ANOVA p<0.001), CTIP2 (**P**, n=4, ANOVA p<0.001), and EdU positive cells distance from the VS (**T**, n=4, ANOVA p<0.001).

To further understand the role of *Piga* in the forebrain and why survival may be compromised, we examined the morphology and myelination of *Piga* mutant brains. Interestingly, the forebrain area in both *Emx1*^*Cre*^; *Piga*^*flox/X*^ females (n=7) and *Emx1*^*Cre/wt*^; *Piga*^*flox/Y*^ males (n=4) did not change compared to wildtype (n=6) at P14 (Fig.S1D-F). Using immuno-staining for myelin basic protein (MBP), we observed moderate hypomyelination in the corpus callosum in both *Emx1*^*Cre/wt*^; *Piga*^*flox/X*^ females and *Emx1*^*Cre/wt*^; *Piga*^*flox/Y*^ males at P14 (SFig.1G-I). *Emx1*^*Cre/wt*^; *Piga*^*flox/X*^ females (n=6) at P21 exhibited a slightly larger cortical area than wildtype (n=12) and *Emx1*^*Cre/wt*^; *Piga*^*flox/Y*^ males (n=9, ANOVA p<0.001, Fig.1B-E). We observed severe hypomyelination in *Emx1*^*Cre/wt*^; *Piga*^*flox/Y*^ males at P21 (Fig.1F-H). Overall, these results show that there is not a consistent effect of *Piga* reduction on dorsal forebrain size and no recapitulation of the microcephaly phenotype as seen in some *PIGA* patients. Myelination deficits seen in the corpus callosum are present in *Emx1*^*Cre/wt*^; *Piga*^*flox/Y*^ animals consistent with findings in male *PIGA* patients.

Prevalent phenotypes in patients with *PIGA* variants include intellectual disability and developmental delay. An explanation for these disabilities is a disorganization of cortical neurons (48), so we proceeded to investigate if *Piga* is required for proper cortical lamination. We examined the upper layer and lower layer neurons in the cortex marked by CUX1 (layers II-III) and CTIP2 (layer V), respectively, at P14 and P21. We observed that upper layer neurons were, on average, found slightly closer to the ventricular surface (VS) in *Emx1*^*Cre/wt*^; *Piga*^*flox/X*^ females (n=4) and *Emx1*^*Cre/wt*^; *Piga*^*flox/Y*^ males (n=4) cortices as compared to wild-type mice (P14, ANOVA p<0.001, SFig.2J-M and P21, ANOVA p<0.001, Fig.1I-L). In addition to calculating the average distance from the ventricular surface, we examined the relative distribution of both specific neuronal subtypes across the cortex. At both P14 and P21, CUX1 positive cells were reduced in upper portions of the cortex (Bin 10) and increased in lower portions (Bins 7,8) in mutant mouse cortices (SFig2.A-B). Conversely, we observed lower layer neurons at slightly further distances from the ventricular surface in *Emx1*^*Cre/wt*^*;Piga*^*flox/X*^ females (n=4) and *Emx1*^*Cre/wt*^; *Piga*^*flox/Y*^ males (n=4) than wild-type mouse brains (n=4, P14 ANOVA p<0.001, SFig.1N-Q and P21 ANOVA p<0.001, Fig.1M-P). We found that CTIP2 positive cells were reduced in Bin 1 and 2 (lower portions of the cortex) and increased in more dorsal regions (Bin 6-7) in mutant mouse cortices (SFig.2C-D). These results show that deletion of *Piga* from the forebrain causes changes in the distributions of both upper- and lower-layer cortical neurons.

We further assessed the *Piga* deletion for effects on radial cell migration using a 5-ethynyl-2’-deoxyuridine (EdU) pulse chase experiment. Pregnant dams producing both *Emx1*^*Cre/wt*^; *Piga*^*flox/X*^ and *Emx1*^*Cre/wt*^; *Piga*^*flox/Y*^ mice were injected with 20 mg/kg of EdU intraperitoneally. EdU then incorporates into the DNA of dividing neural progenitors born in E14.5 embryos, at the height of embryonic upper layer neurogenesis in mice. The brains were collected at P0 and stained for EdU positive cells to observe radial migration of neurons from their birthplace in the ventricular zone to their ultimate destination in the cortical plate. We observed that EdU positive cells were closer to the ventricular surface on average in *Emx1*^*Cre/wt*^; *Piga*^*flox/X*^ females (n=2, p<0.001) than wildtype (n=4,) and *Emx1*^*Cre/wt*^; *Piga*^*flox/Y*^ males (n=4, p=0.171, Fig.1Q-T). Furthermore, *Emx1*^*Cre*^*;Piga*^*flox/X*^ EdU positive cells are distributed differently across the cortex with an increase in EdU positive cells abnormally accumulating in Bins 3 and 5 (SFig.2E). Overall, the results of the cortical marker immunohistochemistry and EdU pulse chase labeling show that reduced levels of *Piga* can adversely affect neuronal radial migration and ultimately cause changes in cortical lamination that could lead to the intellectual disability and developmental delay described in *PIGA* variant patients.

### Piga ablation in oligodendrocytes causes hypomyelination

Delayed myelination and hypomyelination are also common phenotypes of patients with pathogenic *PIGA* variants. This can lead to significant symptoms such as reduced coordination and seizures. The *Nestin-Cre; Piga* model recapitulated the hypomyelination phenotype at P21 in females (41, 49). Oligodendrocytes are a specialized type of brain cell known to deposit myelin in the central nervous system (50-53) and RNA-sequencing data revealed enriched expression of *Piga* in multiple lineages of oligodendrocytes (41, 54). We therefore sought to understand the consequences for *Piga* deficiency in just this population of cells given the *Nestin-Cre; Piga* phenotype. We conditionally ablated *Piga* in the myelinating oligodendrocytes using an *Olig2-Cre* mouse model in which recombination occurs at E17 (50), by producing *Olig2*^*Cre/wt*^; *Piga*^*flox/X*^ female and *Olig2*^*Cre/wt*^; *Piga*^*flox/Y*^ male animals. We first observed the survival at P21 of *Olig2*^*Cre/wt*^; *Piga*^*flox/X*^ females and *Olig2*^*Cre/wt*^; *Piga*^*flox/Y*^ male animals and found that both mutant genotypes survived in close to expected Mendelian ratios (Chi-square p = 0.609; Fig.2A). This revealed that *Piga* ablation in oligodendrocytes alone does not cause the substantial neonatal lethality seen in the *Nestin-Cre; Piga* model. We then observed the extent of myelination using MBP immunohistochemistry in *Olig2-Cre; Piga* mutants. At 3 months of age, we saw reduced levels of MBP along the major axonal tracks into the striatum and the corpus callosum in only *Olig2*^*Cre/wt*^; *Piga*^*flox/Y*^ male brains (black arrows, Fig.2D) while *Olig2*^*Cre/wt*^; *Piga*^*flox/X*^ females (Fig.2C) were unaffected (Fig.2B-D). Interestingly, we did not observe hypomyelination in *Olig2*^*Cre/wt*^; *Piga*^*flox/Y*^ mouse brains at P21 (n=3 animals per genotype, SFig.3A-B), revealing that hypomyelination is a phenotype appearing later in brain development.

**Figure 2.**
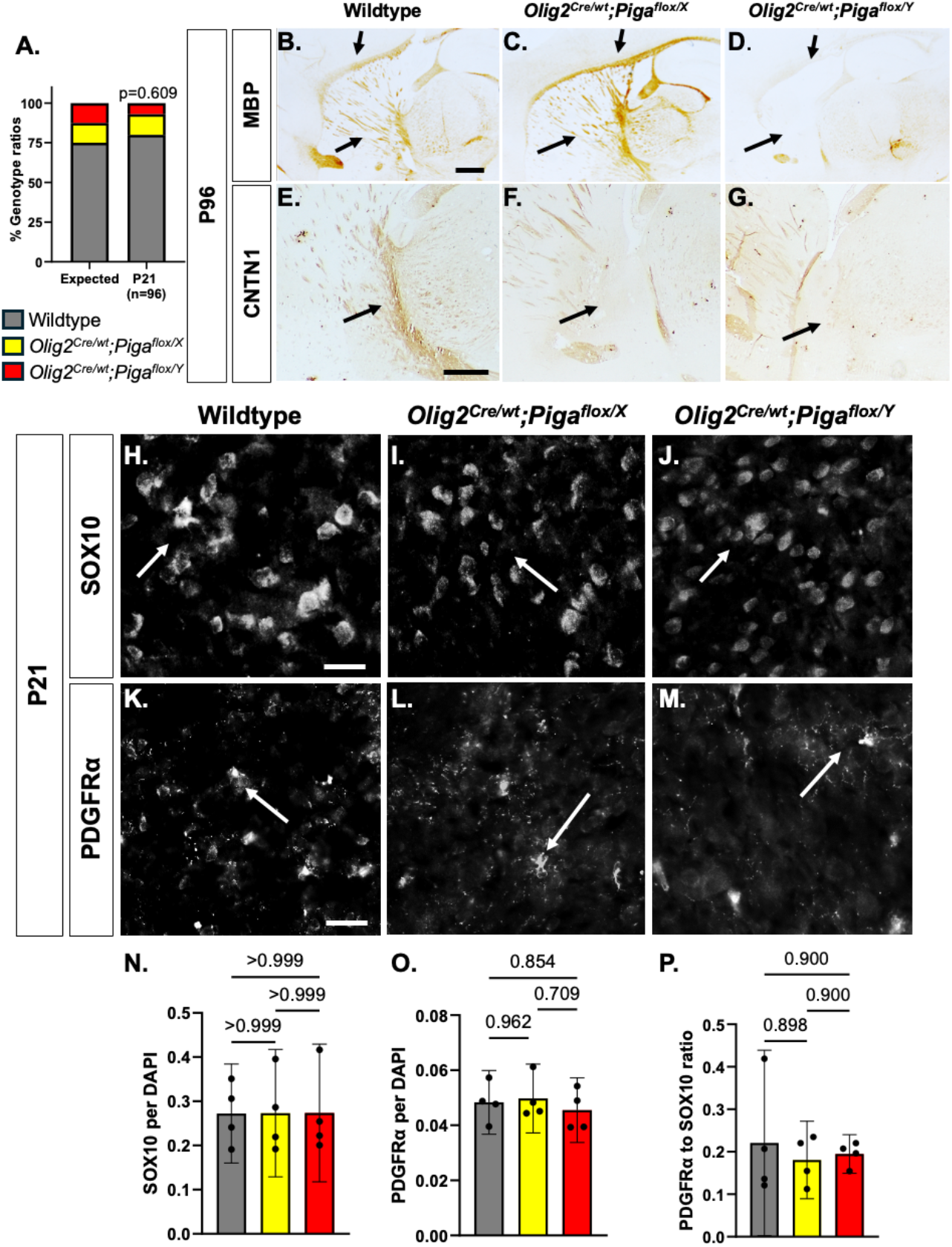
Late onset hypomyelination in *Olig2; Piga* mutant brains. **A)** Percent genotype ratios of wildtype, *Olig2*^*Cre/wt*^; *Piga*^*flox/X*^ female and *Olig2*^*Cre/wt*^; *Piga*^*flox/Y*^ male survival to P21 (n=96, Chi-square p=0.609). Immunostaining of MBP **(B-D)**, CONTACTIN1 **(E-G)**, SOX10 (**H-J**) and PDGFRα (**K-M**) in wildtype (**B**,**E**,**H**,**K**) *Olig2*^*Cre/wt*^; *Piga*^*flox/X*^ female (**C**,**F**,**I.L**) and *Olig2*^*Cre/wt*^; *Piga*^*flox/Y*^ male (**D**,**GJ**,**M**) animals. MBP and CONTACTIN1 data are from P96 (scale bar= 500μm) while SOX10 and PDGFRα are from P21 (scale bar = 25 μm) (n=4 animals per genotype). Quantification of SOX10 (**N**, n=3 animals per genotype, ANOVA p=0.170), PGDFRα (**O**, n=4, ANOVA p=0.564) positive cells and the PDGFR/SOX10 ratio (**P**, n=3, ANOVA p=0.430).

We wanted to further understand why hypomyelination occurs in *Olig2*^*Cre/wt*^*;Piga*^*flox/Y*^ mice. CONTACTIN1 is a GPI-anchored protein and adhesion molecule connecting dendrites of mature oligodendrocytes depositing myelin onto neighboring neuronal axons (53, 55-60). We hypothesize that the loss of *Piga* reduces CONTACTIN1 levels and therefore leads to the hypomyelination phenotype. We used CONTACTIN1 immunohistochemistry and saw that *Olig2*^*Cre/wt*^; *Piga*^*flox/Y*^ male and *Olig2*^*Cre/wt*^; *Piga*^*flox/X*^ female brains (Fig.2F-G) had reduced CONTACTIN1 at 3 months, but not at P21 (Fig.2E-G, SFig.3C-D). These results reveal that *Piga* expression in oligodendrocytes is required for long-term maintenance of myelination and the CONTACTIN1 GPI-anchored protein.

Oligodendrocytes undergo several differentiation stages from oligodendrocyte progenitor cells to myelinating oligodendrocytes (45, 51, 52, 61). Since we observe reduced oligodendrocytes prior to the hypomyelination phenotype in *Olig2*^*Cre/wt*^; *Piga*^*flox/Y*^ males(50-52, 62). SOX10 positive cells in *Olig2*^*Crewt*^; *Piga*^*flox/X*^ females (Fig.2I) and *Olig2*^*Cre/wt*^; *Piga*^*flox/Y*^ males (Fig.2J) were comparable in number to wildtype (Fig.2H) animals at P21 (n=3, ANOVA p=0.170, Fig.2N). We then observed that PDGFRα positive cells are also unchanged in *Olig2*^*Cre/wt*^; *Piga*^*flox/X*^ female (Fig.2L) and *Olig2*^*Cre*^; *Piga*^*flox/Y*^ male (Fig.2M) brains sections relative to wildtype (Fig.2K) animals at P21 (Fig.2K-M, O, n=3 for each genotype, ANOVA p=0.564). Lastly, we observed that the proportion of PDGFRα positive cells to SOX10 positive cells in *Olig2*^*Cre/wt*^; *Piga*^*flox/X*^ and *Olig2*^*Cre/wt*^; *Piga*^*flox/Y*^ animal brains was not markedly different then wildtype animals at P21 (ANOVA p=0.430, Fig.2P). These results show reduced *Piga* expression does not affect the number of mature oligodendrocytes prior to the hypomyelination phenotype observed in the *Olig2*^*Cre/wt*^; *Piga*^*flox/Y*^ mutant males.

### Depletion of Piga from the cerebellum causes early lethality and Purkinje cell disorganization

Cerebellar hypoplasia is one of the brain structural defects caused by human pathogenic *PIGA* variants leading to learning and motor coordination dysfunction (19). We reported the *Nestin*^*Cre/wt*^; *Piga*^*flox/X*^ females exhibited cerebellar hypoplasia and abnormal Purkinje cell arborization (41). To further define the role of *Piga* in the cerebellum, we used the *En1-Cre* which mediates recombination in the hindbrain cell lineage by E9 (46, 63, 64). We crossed *En1-Cre* males with *Piga* flox females to produce *En1*^*Cre/wt*^; *Piga*^*flox/X*^ female and *En1*^*Cre/wt*^; *Piga*^*flox/Y*^ male animals. We first investigated survival and observed that *En1*^*Cre/wt*^; *Piga*^*flox/Y*^ males were underrepresented in comparison to Mendelian expectations by P7 (Chi-square p=0.011, n=48, Fig. 3A) whereas *En1*^*Cre*^; *Piga*^*flox/X*^ females survived to Mendelian expectations. However, myelin and CONTACTIN1 in *Olig2*^*Cre/wt*^; *Piga*^*flox/Y*^ male brains, we hypothesize that loss of *Piga* leads to increased immature oligodendrocytes in the brain at the expense of more mature myelinating oligodendrocytes. We investigated the proportions of PDGFRα-positive oligodendrocyte progenitor cells relative to the total number of SOX10-positive *En1*^*Cre/wt*^; *Piga*^*flox/Y*^ animals did survive to P0 in normal ratios (Chi-square p=0.411, n=109, Fig. 3A). These results are similar to the *Nestin*^*Cre/wt*^; *Piga*^*flox/Y*^ male lethality at P9. We therefore conclude that *Piga* expression is absolutely required in the cerebellum for postnatal survival in males.

**Figure 3.**
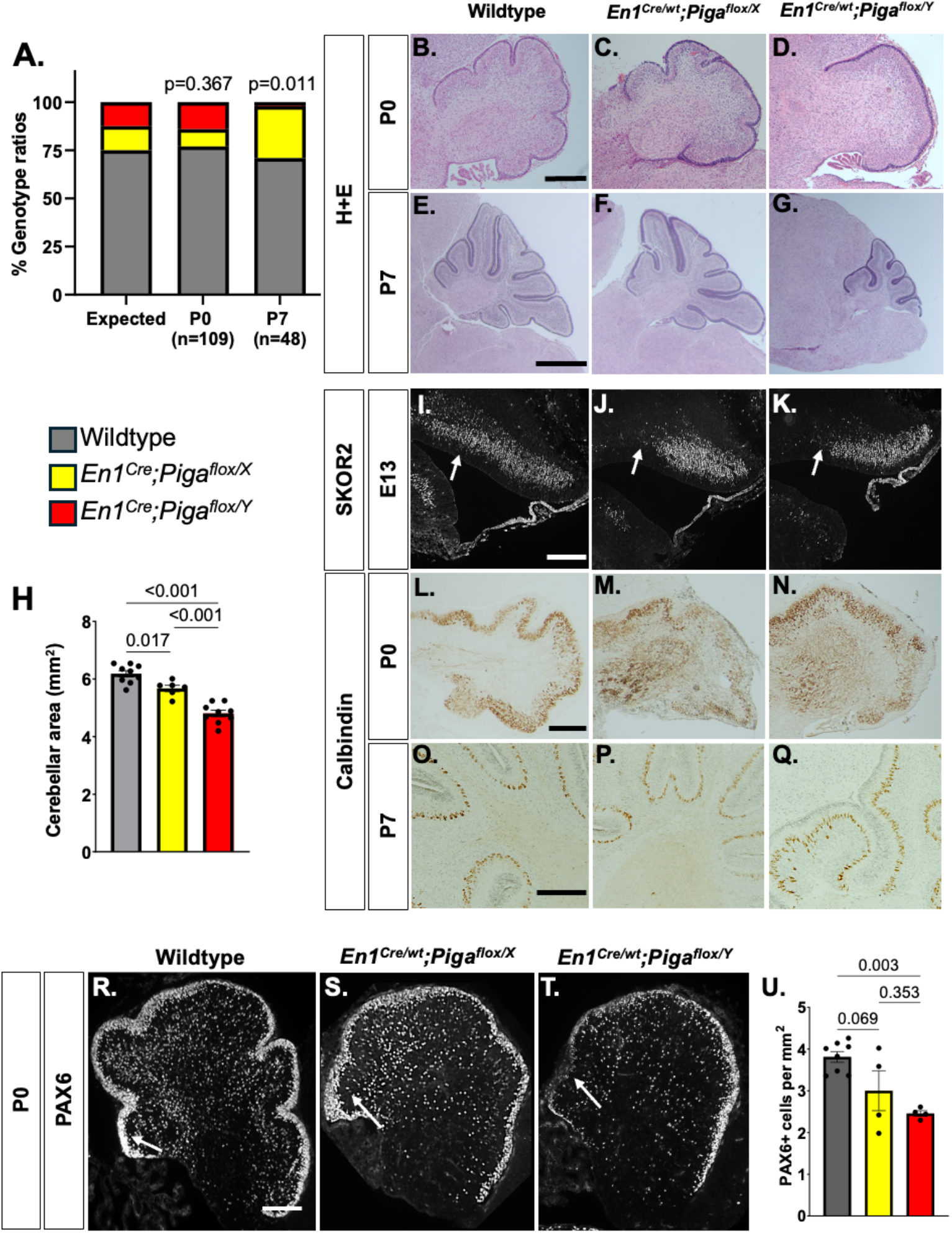
*Piga* is absolutely in the cerebellum required for mammalian survival. **A**) Genotype ratios of *En1*^*Cre/wt*^; *Piga*^*flox/X*^ females and *En1*^*Cre/wt*^; *Piga*^*flox/Y*^ males at P0 (Chi-square p=0.367, n=109) and P7 (Chi-square p=0.011, n=48). H+E stained cerebella (**B-G**) in wildtype (**B, E)** *En1*^*Cre/wt*^; *Piga*^*flox/X*^ females (**C, F**) and *En1*^*Cre/wt*^; *Piga*^*flox/Y*^ males (**D, G**) at P0 (n=4 animals per genotype, scale bars = 250μm) and P7 (1-4 animals per genotype, scale bars= 1 mm). **H)** Quantification of average cerebellar area per animal at P0 (ANOVA p<0.001, 4 sections per animals, n=4 animals each genotype). Immunostaining of SKOR2 marking Purkinje cell progenitors along the ventricular zone in E13 embryo cerebellar hemisphere **(I-K** scale bars= 100 μm), Calbindin marking Purkinje cells (L-Q), and PAX6 marking granule cell progenitors (R-T) in wildtype (**I, L**,**O, R**, n=5-6 animals), *En1*^*Cre/wt*^; *Piga*^*flox/X*^ females (**J, M, P, S**, n=3-6) and *En1*^*Cre/wt*^; *Piga*^*flox/Y*^ males (**K, N, Q, T**, n=3-4 animals, scale bars = 250 μm). **U)** Quantification of PAX6 positive cell per area per animals (average 4 sections per animal). White arrows indicate the anterior domain of the cerebellar ventricular zone at E13. Black arrows point to ectopic levels of Calbindin and white arrows pointing to the posterior domain of the cerebellum at P0.

We analyzed cerebellar growth at P0 when animals of all genotypes could be recovered and observed a reduction in cerebellar area in both *En1*^*Cre/wt*^; *Piga*^*flox/X*^ females (n=4) and *En1*^*Cre/wt*^; *Piga*^*flox/Y*^ males (n=4) animals compared to wildtype (n=4) animals (ANOVA p<0.001, Fig.3B-D, H). We observed also that the single *En1*^*Cre/wt*^; *Piga*^*flox/Y*^ male we obtained at P7 had a significantly smaller cerebellum compared to *En1*^*Cre/wt*^; *Piga*^*flox/X*^ female and wildtype cerebella (Fig.3E-G). We also observed reduced fissures and lobule formation in *En1*^*Cre/wt*^; *Piga*^*flox/X*^ females and *En1*^*Cre/wt*^; *Piga*^*flox/Y*^ males compared to wildtype animal sections at P0, well before the height of cerebellar neurogenesis, to P7 (Fig.3B-G). These results reveal that *Piga* expression is required for early cerebellum morphology and development.

The cerebellar primordium develops as early as E10.5, and at E13.5 Purkinje cell progenitors emerge along the ventricular zone in the cerebellar hemisphere (65-71). Given the cerebellar hypoplasia we noted here and the altered Purkinje cells previously observed (41), we first wanted to observe if *Piga* expression is required for Purkinje cell specification by using the immunohistochemistry against the markers SKOR2 for Purkinje cell progenitors, and Calbindin for Purkinje cells. We see fewer SKOR2 positive cells along the ventricular zone of the cerebellum hemisphere of both *En1*^*Cre/wt*^; *Piga*^*flox/X*^ females (n=6) and *En1*^*Cre/wt*^; *Piga*^*flox/Y*^ males (n=3) compared to wildtype (n=6) sections at E13.5 (Fig.3I-K). Remarkably, the anterior ventricular zone domain (white arrow) was completely lacking SKOR2 positive cells in the mutant mice. At P0, we found increased Calbindin levels in the interior of the cerebellum and a thicker layer of Purkinje cells on the outer surface in *En1*^*Cre/wt*^; *Piga*^*flox/X*^ female (n=5) and *En1*^*Cre/wt*^; *Piga*^*flox/Y*^ males (n=3) animals when compared to wildtype (n=5) animals (Fig.3L-N). At P7, we noticed that the single surviving *En1*^*Cre/wt*^; *Piga*^*flox/Y*^ male had misaligned Purkinje cells (Fig.3O-Q), suggesting a Purkinje cell migration defect. These results reveal that *Piga* expression is required for proper Purkinje cell localization in the cerebellum.

The cerebellar granule cells are the most numerous cells in the central nervous system and develop postnatally in the external granule layer. The underlying Purkinje cells are critical for supporting granule cell proliferation in cerebellar development (72-80). Since Purkinje cells are disrupted in *En1*^*Cre/wt*^; *Piga*^*flox*^ animals, we hypothesized granule cells may also be compromised by a reduction in *Piga*. We observed a reduction in PAX6 positive cells, marking granule cell progenitors, in both *En1*^*Cre/wt*^; *Piga*^*flox/X*^ female (n=4) and *En1*^*Cre/wt*^; *Piga*^*flox/Y*^ male (n=4) cerebella compared to wildtype (n=4) cerebella (ANOVA p=0.003, Fig.3R-U). Overall, these results show that loss of *Piga* has drastic consequences for both Purkinje cell progenitors, even at early embryonic stages, and granule cell progenitors postnatally. We next examined proliferation and cell death, using PHH3 and CC3 antibodies respectively, at P0. We noticed no changes in the number of PHH3 positive cells (ANOVA p=0.582, SFig.4A-D) or CC3 positive cells (ANOVA p=0.378, SFig.4E-H) in *En1*^*Cre/wt*^; *Piga*^*flox/X*^ female and *En1*^*Cre/wt*^; *Piga*^*flox/Y*^ male cerebella compared to wildtype cerebella. Overall, these results reveal that the loss of granule cell progenitors and disrupted Purkinje cell development early in cerebellar development is not due to abnormal cell death or proliferation.

### Ablation of Pgap2 in the forebrain and cerebellum does not compromise survival or brain morphology

Humans with pathogenic *PGAP2* variants exhibit some similar phenotypes to those with loss of *PIGA* including developmental delay, seizures, and hypotonia. However, reports show that *PGAP2* variants lead to less infant lethality and structural brain defects than *PIGA* variants (9, 12, 26, 31, 36, 38, 39). The role of PGAP2 in neural development and the mechanism of such different outcomes has not been previously examined. We have previously reported distinct *Pgap2* expression in the developing mouse forebrain during neurogenesis using a *Pgap2* gene trap mouse allele LacZ reporter (*Pgap2*^*tm1a(EUCOMM)Wtsi*^, SFig.5A). Specifically, we saw expression in the region where intermediate progenitors reside in the developing cortex. Given the importance of these cells to cortical development, we wanted to investigate the role of *Pgap2* in the forebrain (SFig.5B). In addition to reports showing expression in the embryonic cortex, *Pgap2* is highly expressed in the forebrain and cerebellar Purkinje cells at P21 (Fig.4A-B, SFig. 5C-D) (36). This expression pattern is consistent with the intellectual disability and motor dysfunction seen in *PGAP2* patients. To complement the *Piga* analysis presented above, we sought to uncover the role of *Pgap2* in the developing brain and again used *Emx1*-Cre and *En1*-*Cre*. First, we crossed the *Pgap2*^*tm1a*^ conditional gene trap allele or null (81) with *Emx1*-Cre to create *Emx1*^*Cre/wt*^; *Pgap2*^*null*^ animals. We then crossed *Emx1*^*Cre/wt*^; *Pgap2*^*null*^ with *Pgap2*^*tm1c*^ ( or ^flox^) animals to produce *Emx1*^*Cre/wt*^; *Pgap2*^*flox/null*^ mutant animals. We observed that *Emx1*^*Cre/wt*^; *Pgap2*^*flox/null*^ animals survived at Mendelian ratios at P28 (Chi-square test p=0.463, Fig.4C), revealing that *Pgap2* expression in the forebrain is dispensable for survival. We then observed the gross morphology of the mutant *Emx1*^*Cre/wt*^; *Pgap2*^*flox/null*^ compared to wildtype and heterozygous *Emx1*^*Cre/wt*^; *Pgap2*^*flox/wt*^ animals and saw no difference at P28 (Fig. 4D-F). We also saw that body weights of mutant *Emx1*^*Cre/wt*^; *Pgap2*^*flox/null*^ animals (n=3) were comparable to wildtype (n=9) and heterozygous *Emx1*^*Cre/wt*^; *Pgap2*^*flox/wt*^ (n=6) animals (ANOVA p=0.486, Fig.4G). Brain weight was also similar between mutant *Emx1*^*Cre/wt*^; *Pgap2*^*flox/null*^, wildtype, and heterozygous *Emx1*^*Crewt*^; *Pgap2* ^*flox/wt*^ mice (ANOVA p=0.823, Fig.4H) even when normalized to body weight (ANOVA p=0.823, Fig.4H. These results reveal that *Pgap2* in the forebrain is not required for global gross forebrain development. Remarkably, there is a clear difference in brain development between the requirement for GPI-anchor biosynthesis and *Piga* function and the anchor remodeling activity of *Pgap2*.

**Figure 4.**
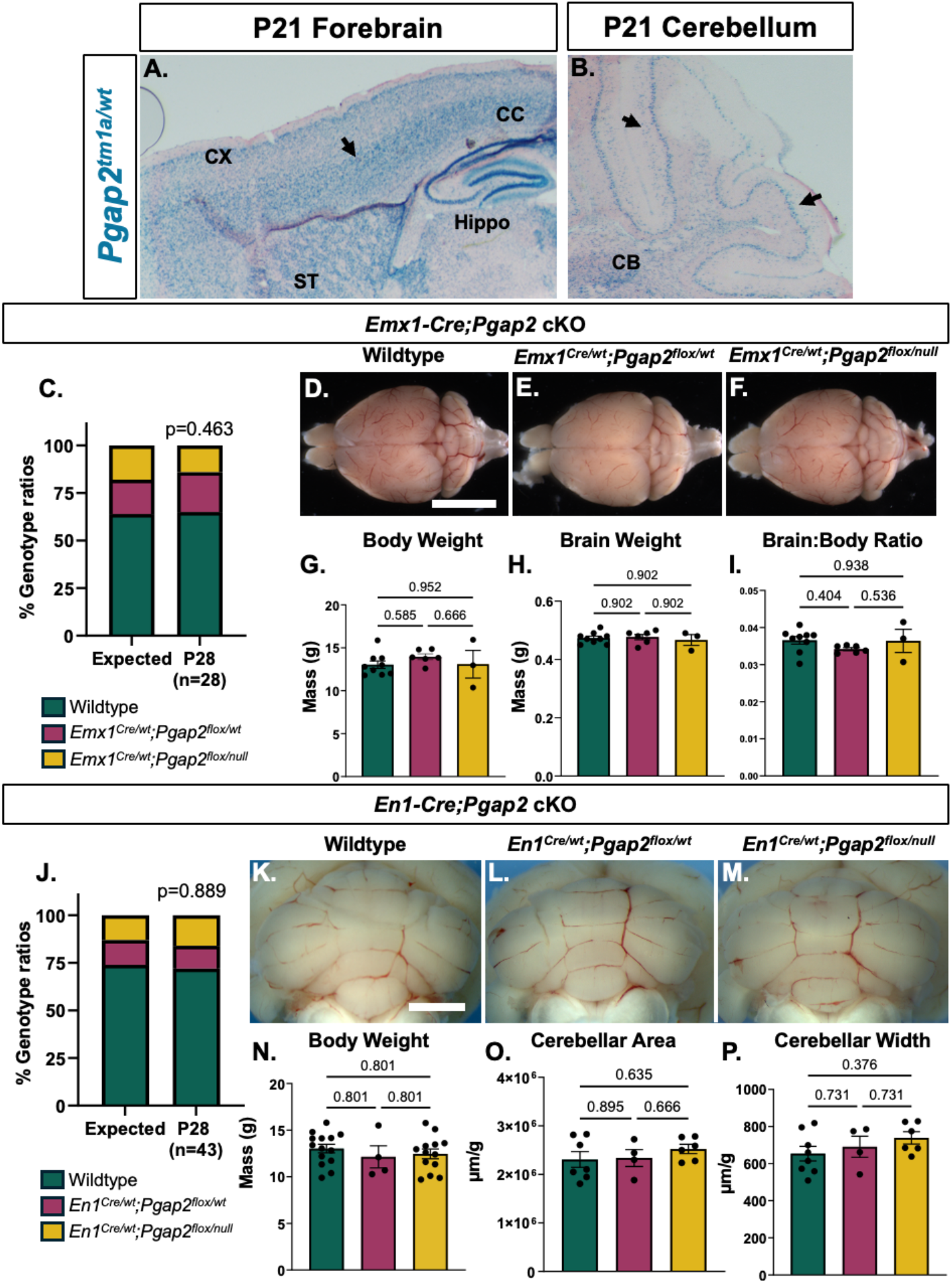
*Pgap2* ablation in the forebrain and the cerebellum does not disrupt brain development. **A)** X-gal stain of *Pgap2*^*null/wt*^ P21 brain sections in the forebrain cortex (CX), striatum (ST), corpus callosum (CC), and **B)** cerebellum (CB). Black arrows pointing to the region of enhanced expression. **C)** Genotype ratios of forebrain ablation *Pgap2* animals at P28 (*Emx1*^*Cre/wt*^; *Pgap2*^*flox/wt*^ heterozygous and *Emx1*^*Cre/wt*^; *Pgap2*^*tm1c/tm1a*^ mutant, n=28, Chi-square p=0.463). **D-F)** Dorsal gross morphological images of wildtype, *Emx1*^*Cre/wt*^*;Pgap2*^*flox/wt*^ heterozygous, and *Emx1*^*Cre/wt*^; *Pgap2*^*null/flox*^ mutants (scale bar = 1 mm). **G-I)** Measurements of body weights, brain weights, and brain to body weight ratio in wildtype, *Emx1*^*Cre*^*;Pgap2*^*flox/wt*^ heterozygous and *Emx1*^*Cre*^; *Pgap2*^*null/flox*^ mutant (n=3-9 animals per genotype). **J)** Genotype ratios of cerebellar ablation of *Pgap2* animals at P28 (*En1*^*Cre/wt*^*;Pgap2*^*tm1c/wt*^ heterozygous and *En1*^*Cre/wt*^; *Pgap2*^*tm1c/tm1a*^ mutant, Chi-square p=0.889). **K-M)** Caudal gross morphological images of the cerebellum in wildtype, *En1*^*Cre*^*;Pgap2*^*flox/wt*^ heterozygous, and *En1*^*Cre/wt*^; *Pgap2*^*null/flox*^ mutants. **N-P)** Measurements of body weights, cerebellar area, and cerebellar width in wildtype, *En1*^*Cre/wt*^*;Pgap2*^*tm1c/wt*^ heterozygous, and *En1*^*Cre/wt*^; *Pgap2*^*tm1c/tm1a*^ mutants (n=4-14 animals per genotype).

We next examined the role of *Pgap2* in the cerebellum. *Pgap2* has distinct expression in Purkinje cells and deletion of *Pgap2* could have a role in survival and cerebellar morphology much like the cerebellar *Piga* mutants. To do this, we crossed the *Pgap2*^*null*^ with *En1*-Cre to create *En1*^*Crewt*^; *Pgap2*^*null/wt*^ animals and crossed those *En1*^*Cre/wt*^; *Pgap2*^*nul/wt*^ animals with *Pgap2*^*flox*^ animals to produce *En1*^*Cre/wt*^; *Pgap2*^*flox/null*^ mutant animals. Mutant *En1*^*Cre/wt*^; *Pgap2*^*flox/null*^ animals survived in Mendelian ratios at P28 (Chi-square test p=0.889, Fig. 4J), revealing that *Pgap2* in the cerebellum is not required for survival, contrary to the absolute requirement for *Piga* in the cerebellum. Since *Pgap2* mutants can survive to P28, we compared the body weight of wildtype, heterozygous *En1*^*Cre/wt*^; *Pgap2*^*flox/wt*^, and mutant *En1*^*Cre/wt*^; *Pgap2*^*flox/null*^ animals at P28 and found that there was no significant difference in body weight (ANOVA p=0.614, Fig. 4N). We then measured cerebellar area, height and width and vermis height and width relative to body weight to quantify the effect of deletion of *Pgap2* on cerebellar morphology (SFig.5E-H, Figure 4K-M). We observed no significant difference in vermis width (ANOVA p=0.502, Fig.S5F), vermis height (ANOVA p=0.643, Fig. S5G), cerebellum height (ANOVA p=0.601, SFig.5H), cerebellar area (ANOVA p=0.523, Fig.4O) or cerebellar width (ANOVA p=0.335, Fig.4P) between mutant *En1*^*Cre/wt*^; *Pgap2*^*flox/null*^, heterozygous *En1*^*Cre/wt*^*;Pgap2*^*flox/wt*^, and wildtype animals. Overall, these results show that *Pgap2* is not required in the forebrain and cerebellum for survival and brain morphology by P28 in mouse models. These results also suggests that *Piga* and *Pgap2*, though in the same GPI-anchor biosynthetic pathway, have vastly different requirements in the developing brain.

## Discussion

Inherited GPI deficiencies can cause many congenital anomalies with the most prevalent being intellectual and developmental delay, and structural brain anomalies (11, 13, 26, 82). Specifically, *PIGA* patients exhibit hypomyelination, cerebellar hypoplasia, some cases of microcephaly and most devastatingly, infant mortality (11, 18, 19, 22-25, 27-29, 33, 83). Our previous studies with a central nervous system specific deletion of *Piga* (*Nestin-Cre*) in mice recapitulated many human symptoms suggesting this is a valid model for these *PIGA* human variants(41). *Pgap2*, however, does not have any mouse models to study its role in the brain. In this study, we provide major findings: 1) *Piga* expression is required most acutely in the cerebellum, and 2) *Piga* and *Pgap2* have dramatically different requirements in neural development.

The phenotypes from the *Olig2*-*Cre* and *Emx1-Cre* ablations of *Piga* from oligodendrocytes and the dorsal forebrain did not manifest until after weaning age which was outside the scope of these neurodevelopment studies. The common and most severe phenotype in the *Olig2-Cre* and *Emx1-Cre* deletions was hypomyelination. However, the hypomyelination occurred early in the *Emx1-Cre;Piga* mutants and later in *Olig2-Cre;Piga* mutants. This could be due to the timing of recombination as *Emx1-Cre* is active earlier at E10.5 as compared to the *Olig2-cre* which begins expression at E17, well after neurogenesis begins (45). Despite the subtle phenotypes, there are opportunities to investigate further why *Olig2-Cre;Piga* animals have late onset hypomyelination. Myelin turns over in the central nervous system and new mature oligodendrocytes must be produced (50-52, 60). It is still unclear if *Piga* has a role in the regeneration of mature oligodendrocytes. However, it is clear from our work that *Piga* depletion from oligodendrocytes does not cause the early lethality seen in *Nestin-Cre; Piga* mutants. More long-term studies will need to be performed on *Olig2-Cre;Piga* animals to determine the role of *Piga* in oligodendrocyte development.

We were surprised to see only subtle cortical defects in *Emx1-Cre; Piga* mutant forebrain. A previous report showed that the male *Emx1-Cre; Piga* mouse model died embryonically due forebrain abnormalities (19). The *Emx1-Cre* allele used in that study was not the more generally used reagent we employed here (84). The death seen in the previous report was both earlier than one would predict from a forebrain phenotype, and difficult to classify as true microcephaly given overall tissue necrosis. The *Emx1*-Cre; *Piga* male mice presented here die right after weaning for reasons that are not yet understood. Those mutant mice have hypomyelination in the corpus callosum which is known to cause seizures in patients (18, 28). We hypothesize this localized hypomyelination may lead to seizure activity that is lethal in adult mice, but further studies are needed to test this hypothesis.

We found that deletion of *Piga* from the hindbrain (*En1-Cre; Piga*) had the largest effect causing early lethality and aberrant cell morphology. First, we have attempted to observe cerebellar morphological changes in *En1*^*Cre/wt*^; *Piga*^*flox/Y*^ males at P7 when the external granule layer forms which is one of most important steps in cerebellar morphogenesis. However, the majority of *En1*^*Cre/wt*^; *Piga*^*flox/Y*^ males die between P0 and P5 so it is very challenging to robustly analyze how *Piga* expression affects cerebellum development. Overall, we were still able to provide new evidence for *Piga*’s role in Purkinje cell specification and localization at stages up to birth. In addition to the disruptions to Purkinje cells and granule cells we see which may affect motor coordination, other hindbrain functions including cardiovascular and respiratory control may lead to this early lethality ^12,134^. These questions may be fruitful avenues for future investigation. Nonetheless, in the context of understanding treatment options, our results suggest that the cerebellum is a potential target tissue for therapies to treat *PIGA*-related disorders.

We observed that *Pgap2* deletion did not affect brain structure like the deletion of *Piga*, even though these enzymes are in the same GPI-anchor biosynthesis pathway. We do note that patients with *PGAP2* pathogenic variants exhibit less severe symptoms than *PIGA* patients (35). We hypothesize these results could be due to their sequence of activity within the GPI-anchor biosynthesis cascade. PIGA is one of the first enzymes in the pathway, acting in the endoplasmic reticulum, whereas PGAP2 is one of the last enzymes in the pathway in the Golgi, after proteins are bound to the anchor, with a role in remodeling the anchor. Our data suggest that in neural development, remodeling of the GPI-anchor is much less critical than the initial creation of the anchor. Altogether, we have expanded our knowledge of the role of GPI-anchor biosynthesis enzymes in several neuronal cell types.

## Acknowledgments

We would like to thank the Stottmann laboratory members for their continuous council throughout this study. Specifically, we thank Anthony Koulianos for his part in data collection. We would like to thank Dr. Waclaw for his discussion about oligodendrocyte biology. Lastly, we would like to thank the funding sources from Nationwide Children’s Hospital Recruitment Funds and the Abigail Wexner Research Institute Postdoctoral Diversity in Academia award.

## Author contributions

**Conceptualization**: R.W.S., J.L.W; **Methodology:** J.L.W., R.W.S..; **Validation:** J.L.W., J.M.L. R.W.S.; **Formal analysis**: J.L.W.; **Investigation**: J.L.W., J.M.L.; **Resources**: R.W.S. **Writing - original draft:** J.L.W; **Writing - review & editing:** J.L.W., J.M.L., R.W.S; **Visualization:** J.L.W; **Supervision**: R.W.S.; **Project administration:** R.W.S.; **Funding acquisition**: R.W.S., J.L.W.

## RESOURCE AVAILABILITY

### Lead contact

Further information and requests for resources and reagents should be directed to and will be fulfilled by the lead contact, Rolf Stottmann

### Materials availability

Materials will be fulfilled by the lead contact upon request

### Data and code availability

All data in this paper will be shared by the lead contact upon request.

No new code was produced in this paper Any additional information about the data acquired in this paper for further analysis is available from the lead contact upon request.

## Materials and Methods

### Animal husbandry

All animals were maintained through a protocol approved by the Nationwide Children’s Hospital IACUC committee (IACUC: AR21-00067). Mice were housed in a vivarium with a 12-hour light cycle with food and water *ad libitum*. A series of mice in the study were obtain through multiple repositories with their genotyping scheme shown in methods table. *Piga*^*flox*^ (*B6*.*129-Piga*^*tm1*^, #RBRC06211)(40) mice were previously generated by Taroh Kinoshita and Junji Takeda and obtained from the RIKEN repository. Mice were genotyped for the *Piga*^*flox*^ allele as published (Methods Table 1). *B6*.*129-Olig2*^*tm1*.*1(cre)Wdr/J*^*or Olig2-Cre* (Jackson labs #025567) (45), *B6*.*129S2-Emx1*^*tm1(cre)Krj/J*^ or *Emx1-Cre* (Jackson labs #005628) (44) animals were genotyped using Jackson Lab’s recommended general Cre genotyping. *En1*^*tm2(cre)Wrst/J En1-Cre*^ or *En1-Cre* animals (Jackson labs #007916) (46) were genotyped using the *En1-Cre* specific primer from Jackson Labs. Individual Cre and flox mouse lines were maintained on a C57Bl6/J background. Progeny of the *Piga*^*flox*^ and Cre crosses were euthanized at the proper humane endpoint.

**Table 1.**
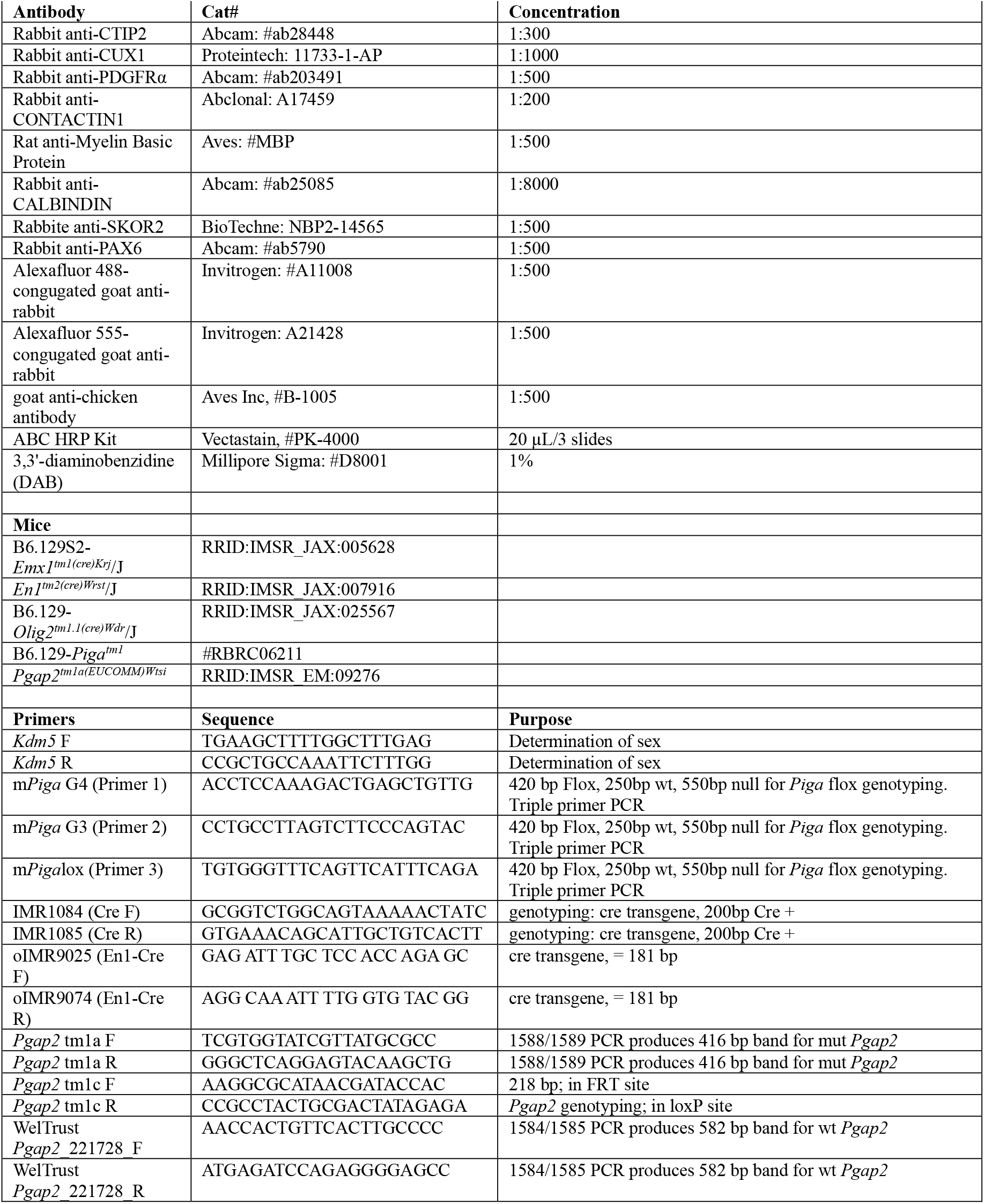
Reagents.

### Histology and Immunostaining

Brains were harvested and fixed in 10% formalin or 4% paraformaldehyde for 24–48 hr. Brains were washed in either 70% ethanol and paraffin embedded by the NCH Biopathology Core or 30% sucrose and embedded in O.C.T. solution. Brains were sectioned by microtome or cryostat at 10 μm and stained with hematoxylin & eosin.

Immunohistochemistry was performed on formalin-fixed, paraffin embedded brain tissue. Tissue was sectioned at 10 μm, sections were blocked for 1hr at room temperature in 4% normal goat serum in PBST and incubated in primary antibody overnight at 4°C. The next day, slides were washed in PBS and incubated in 1:500 biotinylated secondary antibody for 1hr at room temperature. The slides were washed and incubated in ABC mix (Vectastain ABC HRP Kit, #PK-4000) for 1 hour at room temperature. The slides were washed and developed in 0.5 mg/ml DAB (Sigma) activated with 30% hydrogen peroxide. Slides were incubated in 3,3’-diaminobenzidine (DAB) for approximately 5 min, washed in PBS, sealed with cytoseal and imaged by light microscopy.

Immunofluorescence was performed on PFA fixed, OCT embedded samples. Frozen sections were heated on a heat block at 42°C for 10-15 mins. Antigen retrieval was performed with citrate retrieval or vector commercial buffer, blocked in 4% normal goat serum, and incubated with the following primary antibodies overnight at 4°C. Sections were incubated with secondary antibodies and counterstained with DAPI. Sections were imaged on the Zeiss Apotome Fluorescence Microscope.

### Statistical analysis

Statistical analysis and graphs were generated using GraphPad Prism (GraphPad Software, San Diego, CA). A Chi-square test was used to analyze genotype ratios. Data shown in bar graphs and scatter plots are presented as the median with the Standard Error of the Mean (SEM). Survival analysis was performed with a log-rank Mantel-Cox test. Cell and area quantification analysis were performed with a one-way ANOVA followed by Tukey’s multiple comparison analysis. P-values shown in bar graphs are the results of the multiple comparisons testing unless otherwise specified.

