## Supplemental Information for "Genetic Dissection of the role of *Piga* and *Pgap2* in the embryonic mouse brain"

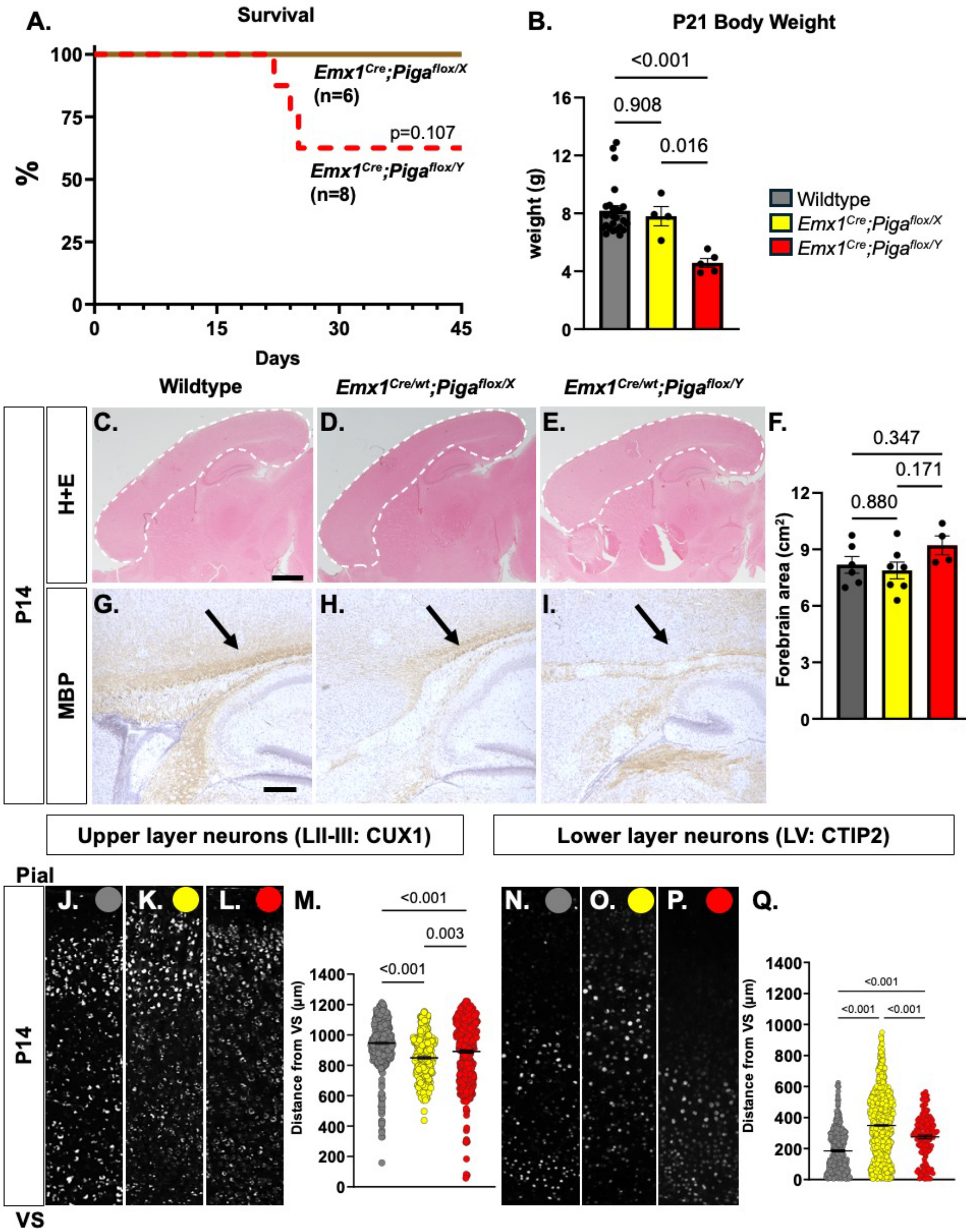


**Figure S1. *Piga* ablation in forebrain reduces survival in mutant males shortly after weaning. A)** Kaplan-Meier plot of survival of *Emx1*$^{Cre/wt}$; *Piga*$^{flox/X}$ female and *Emx1*$^{Cre/wt}$; *Piga*$^{flox/Y}$ male up to P45 (Mantel-Cox p=0.107). **B)** Body weight of wildtype (n=25), *Emx1*$^{Cre/wt}$; *Piga*$^{flox/X}$ females (n=7), and *Emx1*$^{Cre/wt}$; *Piga*$^{flox/Y}$ males (n=5) at P21 (ANOVA p<0.001). **C)**. Representative H+E images of wildtype (n=6) , **D)** *Emx1*$^{Cre/wt}$; *Piga*$^{flox/X}$ female (n=7) and **E)** *Emx1*$^{Cre/wt}$; *Piga*$^{flox/Y}$ male (n=4) P14 forebrains (scale bar= 1000 μm). **F)** Quantification of forebrain area in images C-E (ANOVA: p=0.009). Immunostaining of MBP (**G-I**, scale bar= 250 μm), CUX1 (**J-L**), and CTIP2 (**N-P**) in wildtype (**G, J, N**) *Emx1*$^{Cre/wt}$; *Piga*$^{flox/X}$ female and (**H, K, O**) *Emx1*$^{Cre/wt}$; *Piga*$^{flox/Y}$ male P14 forebrains (**I, L, P**, n=4 animals per genotype). **M)** Quantification of CUX1 (**M,** ANOVA p<0.001) and CTIP2 (**Q**, ANOVA p<0.001) positive cells distance from the VS.

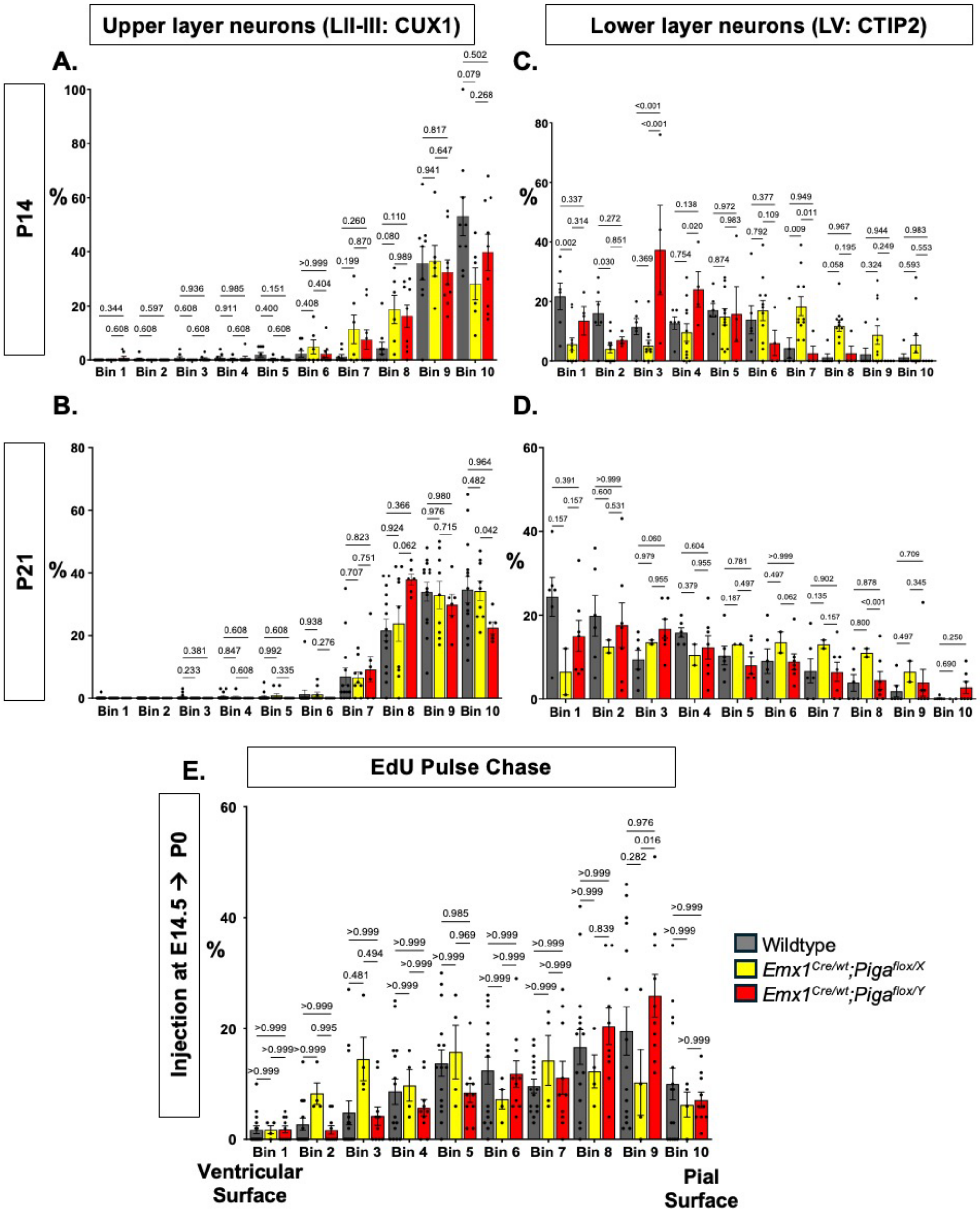


**Figure S2. Distribution of upper and lower layer neurons in *Piga* depleted forebrains.** Percent of Upper layer neurons CUX1 positive cells in wildtype, *Emx1*$^{Cre/wt}$*; Piga*$^{flox/X}$ female and *Emx1*$^{Cre/wt}$*; Piga*$^{flox/Y}$ male forebrains in bin 1 ventricular surface to bin 10 pial surface at **A)** P14 and **B)** P21 and lower layer neurons at **C)** P14 and **D)** P21. **E)** Percent of EdU positive cells in P0 pups forebrains injected at E14 from the bin 1 VS to bin 10 pial (n=2-4 animals per genotype)

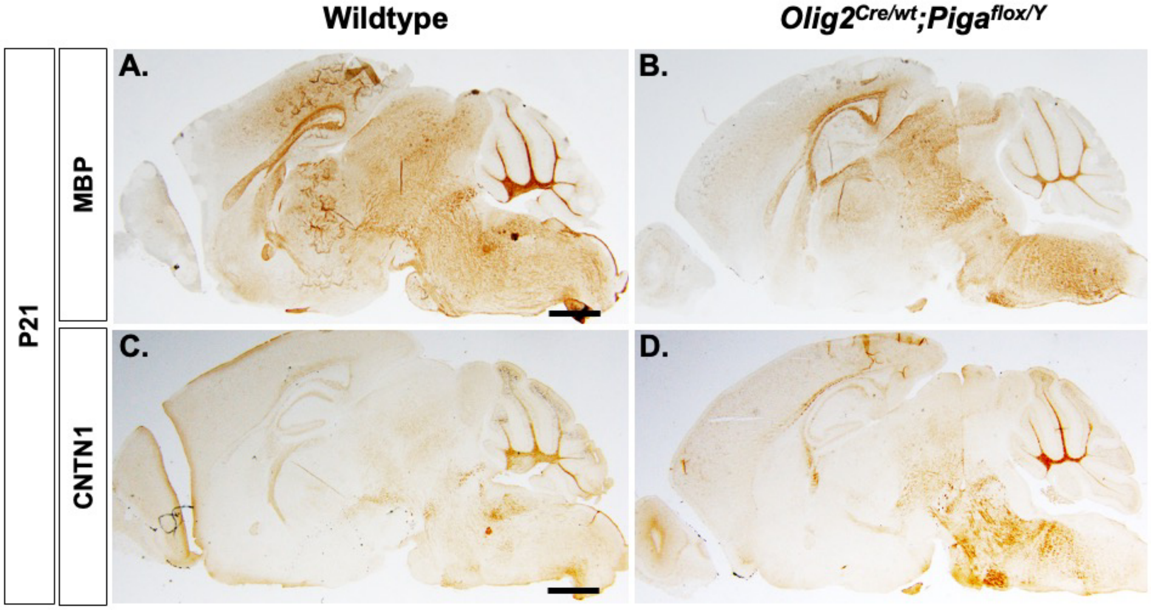


**Figure S3. Myelin and CONTACTIN1 proteins are not disrupted in *Piga* ablated oligodendrocyte males at P21. A)** wildtype and **B)** *Olig2$^{Cre/wt}$; Piga$^{flox/Y}$* male P21 brain sections immunostained with MBP. **C)** Immunostaining of CONTACTIN1 in wildtype and **D)** *Olig2$^{Cre/wt}$; Piga$^{flox/Y}$* male P21 brain sections (n=4 animals per genotype).

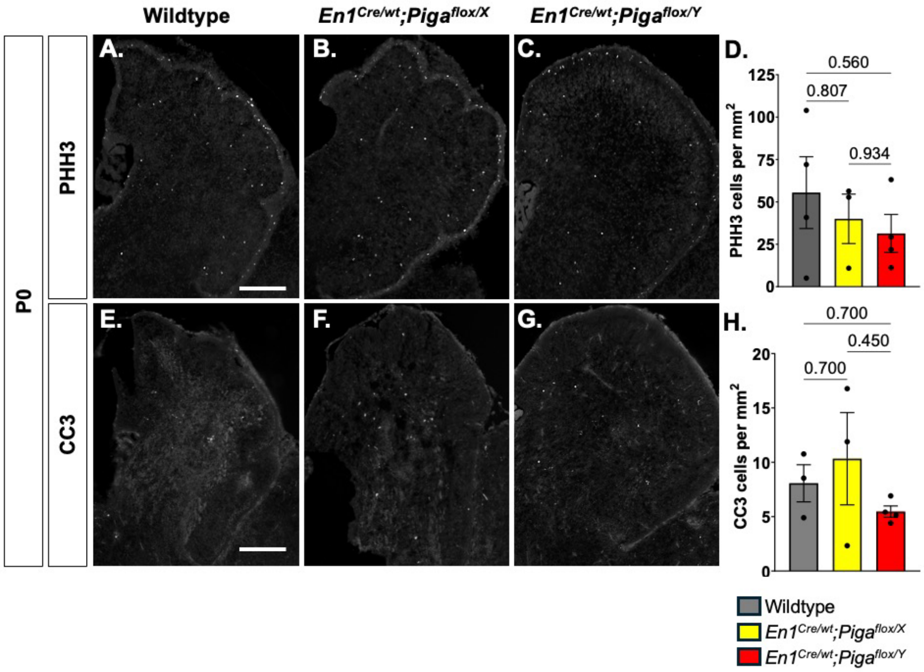


**Figure S4. Cell proliferation and cell death is not disrupted in *Piga* depleted cerebellum cells at P0.** P0 cerebella immunostained with PHH3 marking proliferative cells (**A-C**) and CC3 (**E-G**) marking cell deathin wildtype (**A, E**, n=3-4),*En1$^{Cre/wt}$; Piga$^{flox/X}$* females (**B, F**, n=3) and*En1$^{Cre/wt}$; Piga$^{flox/Y}$* males (**C, G**, n=3-4) cerebella (scale bars= 250 µm). The average PHH3 (**D**) and CC3 (**H**) positive cells are quantified average 4 sections per animal per genotype.

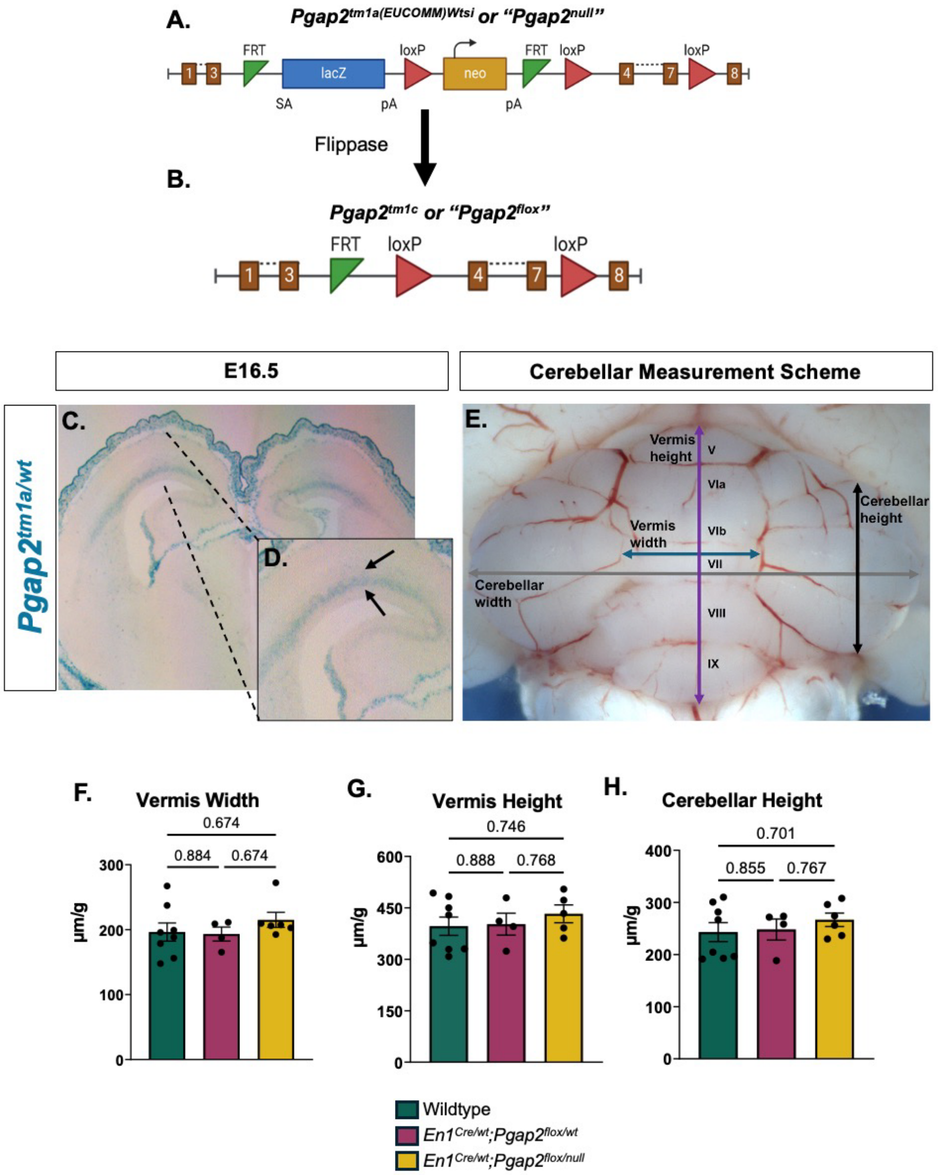


**Figure S5.** ***Pgap2*** **expression in the developing forebrain. A)** Gene trap scheme of $Pgap2^{tm1a}$ and **B)** $Pgap2^{tm1c}$ allele in conditional knockout *Pgap2* experiments. **C)** X-gal stain of *Pgap2* expression in coronal E16.5 embryos. **D)** Zoomed in image of *Pgap2* expression in E16.5 mouse embryo cortex. **E)** Lines depicting cerebellar measurements for cerebellar width, cerebellar height, vermis width, and height. **F)** Measurements of vermis width, **G)** vermis height, and **H)** cerebellar height for wildtype, $En1^{Cre/wt}$; $Pgap2^{tm1c/wt}$ heterozygous, and $En1^{Cre/wt}$; $Pgap2^{tm1c/tm1a}$ mutants.
